# β-arrestin-biased Allosteric Modulator of Neurotensin Receptor 1 Reduces Ethanol Drinking and Responses to Ethanol Administration in Rodents

**DOI:** 10.1101/2024.04.10.588903

**Authors:** Graydon B. Gereau, Diana Zhou, Kalynn Van Voorhies, Ryan E. Tyler, Jeffrey Campbell, Jackson G. Murray, Ali Alvarez-Pamir, Luke A. Wykoff, Michel A. Companion, Michael R. Jackson, Steven H. Olson, Lawrence S. Barak, Lauren M. Slosky, Ryan P. Vetreno, Joyce Besheer, Zoe A. McElligott

## Abstract

Alcohol use disorders (AUDs) impose an enormous societal and financial burden, and world-wide, alcohol misuse is the 7^th^ leading cause of premature death^1^. Despite this, there are currently only 3 FDA approved pharmacological treatments for the treatment of AUDs in the United States. The neurotensin (Nts) system has long been implicated in modulating behaviors associated with alcohol misuse. Recently, a novel compound, SBI-553, that biases the action of Nts receptor 1 (NTSR1) activation, has shown promise in preclinical models of psychostimulant misuse. Here we investigate the efficacy of this compound to alter ethanol-mediated behaviors in a comprehensive battery of experiments assessing ethanol consumption, behavioral responses to ethanol, sensitivity to ethanol, and ethanol metabolism. Additionally, we investigated behavior in avoidance and cognitive assays to monitor potential side effects of SBI-553. We find that SBI-553 reduces binge-like ethanol consumption in mice without altering avoidance behavior or novel object recognition. We also observe sex-dependent differences in physiological responses to sequential ethanol injections in mice. In rats, we show that SBI-553 attenuates sensitivity to the interoceptive effects of ethanol (using a Pavlovian drug discrimination task). Our data suggest that targeting NTSR1 signaling may be promising to attenuate alcohol misuse, and adds to a body of literature that suggests NTSR1 may be a common downstream target involved in the psychoactive effects of multiple reinforcing substances.

## Introduction

### Neurotensin systems and alcohol

Neurotensin (Nts) systems have previously been explored in the context of and in circuitries related to AUD, stress disorders, and motivated behaviors^2–7^. Levels of neurotensin in the cortex are significantly lower in Alcohol Preferring P-rats as compared to their controls^8^. Similarly, mice bred for variance in ethanol sensitivity (Long Sleep/LS and Short Sleep/SS mice), differentially express both neurotensin and neurotensin receptors. Ethanol and neurotensin cross-sensitize to locomotor behaviors, and the actions of each compound on hypothermia responses^9,10^. Ethanol Administration promotes cross-tolerance to the locomotor effects of neurotensin, and i.c.v. administration of neurotensin enhances the sedative properties of ethanol^9,10^ Global knockouts of NTSR1 exhibit enhanced ethanol consumption and preference and reduced ataxia to ethanol (1.5 g/kg i.p.), while not showing psychomotor stimulation or depression by low doses of ethanol (1 – 1.5 g/kg ethanol i.p.)^11^. Finally, human genetic studies have found that polymorphisms in the *Ntsr1* gene are significantly associated with AUD and substance use disorders^12,13^.

Because of these studies, we have previously investigated how the central nucleus of the amygdala (CeA) neurons expressing Nts regulate ethanol consumption and reinforcing behaviors^4^. In male but not female mice, we found that ablation of or knockdown of GABA release from CeA Nts neurons reduced ethanol consumption, while stimulation of the terminal projections to the parabrachial nucleus enhanced the consumption of ethanol and was reinforcing on its own in ethanol naïve mice^2,4^. Furthermore, we have shown that endogenous neurotensin release in the bed nucleus of the stria terminalis (BNST) enhances synaptic transmission between the CeA and BNST, a circuit that promotes ethanol consumption^14,15^. Finally, it is worth noting that alcohol and Nts have an important relationship given the roles of Nts in feeding behavior, mediating metabolic processes, consumption, and psychosis^7,16,17^. Meanwhile, alcohol is a distinctive misused substance in that it is both highly caloric and psychoactive, which makes this neuropeptide/misused substance relationship somewhat unique and complex^7,17^.

### Biased Signaling and SBI-553

G protein coupled receptors, like NTSR1, can be selectively stabilized by ligands in distinct conformational states that bias signaling towards distinct pathways, e.g. some being mediated by specific heterotrimeric G proteins or β-arrestins. Agonists and/or modulators can promote the receptor to adopt a confirmational state that biases the activation of the receptor towards or away from certain intracellular signaling cascades^18^. Recently, it was reported that a novel β-arrestin-biased positive allosteric modulator of NTSR1, SBI-553, increases Nts-NTSR1 occupancy, antagonizes NTSR1-associated Gq activity and potentiates β-arrestin signaling at this receptor when bound by an agonist^19,20^. This small molecule altered animal models of psychostimulant use, without engaging the aversive side effects of other non-biased modulators and/or agonists of neurotensin receptors such as hypotension and hypothermia^19^. Reductions in conditioned place preference (CPP) recall and self-administration of psychostimulants point to strong potential for this compound to engage in modulation of ethanol-related behaviors given the relationship between Nts and ethanol/AUD outlined above^19^.

Because of the aforementioned impact of neurotensin signaling on various aspects of ethanol-related behaviors, here we examine if this potential pharmacotherapy has efficacy in changing ethanol drinking and related measures. We find that systemic SBI-553 treatment alters aspects of ethanol-associated behavior including a reduction in the amount of ethanol consumed in binge-like ethanol drinking models (2-bottle and single bottle drinking in the dark, or DID), but not sucrose consumption in mice. We find that this effect is not related to modulation of avoidance behaviors. We also find that SBI-553 modulates physiological response to ethanol administration. Additionally, drug discrimination experiments in rats demonstrate that SBI-553 attenuates interoceptive effects of ethanol. These findings lead us to hypothesize that reductions in ethanol drinking could be due to modulation of the interoceptive qualities of ethanol by SBI-553. These data further support that the Nts system, and NTSR1 in particular, may be a promising target for the treatment of multiple substance use disorders, and that SBI-553 could potentially be a pharmacotherapeutic for the treatment of AUD.

## Methods

### Animals

All rodents were under continuous care and monitoring by veterinary staff from the Division of Comparative Medicine at UNC-Chapel Hill. All procedures were conducted in accordance with the NIH Guide to Care and Use of Laboratory Animals and UNC Institutional Animal Care and Use Committee approvals and guidelines.

Adult (8 weeks at time of delivery) male and female C57BL6/J (Jackson Laboratories) were used for all experiments expect for drug discrimination experiments in rats. Mice were allowed at least one week to habituate in the vivarium before testing. The vivarium was on a 12hr light/dark reverse light cycle, with experiments conducted during the dark cycle for all experiments. Mice for drinking experiments were individually housed one week prior to drinking. All mice had *ad libitum* access to food and water, with the exception of the novelty induced suppression of feeding test (NSFT, see below).

Adult (7 weeks at the time of delivery) male and female Long Evans rats (Envigo, Indianapolis, IN) were used for Pavlovian ethanol discrimination experiments only. Rats were individually housed. The vivarium was on a 12-hr light/dark cycle and all experiments were conducted during the light cycle. All rats received daily handling for at least 1 min/day for 1 week prior to the start of discrimination training. Food intake was restricted to maintain body weight (males: 320-340 g; females: 230-255 g). Rats had *ad libitum* access to water.

### Treatments

SBI-553 (Sanford Burnham Presbys Medical Discovery Institute, San Diego, CA) was dissolved in 5% hydroxypropyl β cyclodextrin and 0.9% sterile saline at a concentration of 1.2mg/mL SBI-553. Vehicle used in studies for this paper was 5% hydroxypropyl β cyclodextrin and 0.9% sterile saline. Mice received injections of 12mg/kg SBI-553 or an equivalent volume of vehicle i.p. 30 minutes prior to each test. For some avoidance tests, mice were treated with 20% ethanol in 0.9% sterile saline, just before the start of those tests. Rats received injections of 2, 6, or 12mg/kg SBI-553 i.p. or an equivalent volume of vehicle for the Pavlovian ethanol discrimination study, prior to i.g. ethanol or water delivery.

### Drinking in the Dark (DID)

DID was conducted based on previously described methods^21^. Mice were given *ad libitum* access to chow and water in the home cage. At the start of the experiment, one water bottle was replaced by a bottle containing 20% (weight/volume) ethanol or 3% sucrose at 10AM, and was removed at 12 PM. This process is repeated for two additional days before the test day (Day 4). On the test day, mice were injected with 12mg/kg SBI-553 or saline 30 minutes prior to the ethanol bottle placement. We measured drinking in both one-bottle and two-bottle choice DID paradigms in separate cohorts of mice. In two-bottle choice DID the bottle position was rotated across days to eliminate a side preference. Ethanol and water consumption were measured at the end of the test period.

### Elevated Plus Maze (EPM)

Mice were treated with SBI-553 12mg/kg (i.p.) or vehicle 30 minutes prior to a 1g/kg ethanol injection (i.p.). In separate experiments, no ethanol was administered to investigate SBI-553 effects alone. Mice were placed in the center of an elevated plus maze and allowed to explore the maze for 5 minutes. Time spent in the closed arms, open arms, and center of the maze were tracked using Ethovision XT 14 (Noldus).

### Open Field

Mice were treated with SBI-553 12mg/kg (i.p.) or vehicle 30 minutes prior to a 1g/kg ethanol injection (i.p.) or no ethanol injection for separate experiments investigating SBI-553 on its own. Mice were allowed to freely explore a 42cm x 42cm novel open field for 30 minutes. Locomotor data were collected using Fusion software (Omnitech).

### Blood Ethanol Concentration Assay

Mice were treated with SBI-553 12mg/kg (i.p.) or vehicle 30 minutes prior to a 2g/kg ethanol injection (i.p.). 15 minutes after ethanol injection, mice were briefly anesthetized using isoflurane and sacrificed 30 seconds later. Whole trunk blood was collected and analyzed on an Analox AM1.

### Ethanol Sedation Assay

Mice were treated with either 12mg/kg SBI-553 or vehicle (i.p.) 30 minutes prior to receiving a 4g/kg dose of ethanol (i.p.). Following the ethanol injection, a timer was started and mice were placed back into their home cage until they were visibly sedated. Once visibly sedated, mice were placed in the supine position to confirm loss of righting reflex (LORR). At this point mice were moved to individual troughs, eye lubricant was applied, and the time it took to right themselves three times in a 30-seconds window was measured as the LORR measure (righting time – time of sedation).

### Ethanol Response Battery (ERB)

Ethanol response battery was conducted as previously described^22^. In a counterbalanced fashion, mice underwent ERB assessment beginning at 8:00 AM consisting of a baseline assessment after treatment with SBI553 (12mg/kg, i.p.) or vehicle, followed by four consecutive cumulative doses of ethanol to assess ethanol responsivity across a broad range of blood ethanol concentrations (BECs). The four consecutive doses of ethanol (i.e., 0.5, 1.0, 1.0, 1.0 g/kg, i.p.) produce cumulative ethanol doses of approximately 0.5, 1.0, 2.0, and 3.0 g/kg. Each subsequent ERB assessment was conducted approximately 30 min apart, each initiated 15 min after ethanol dosing. The ERB consisted of (1) the 6-point behavioral intoxication rating scale, (2) body temperature assessment, (3) submandibular, (4) tilting plane assessment, and (5) LORR following the final ethanol dose. Body weight was assessed at the beginning of the ERB.

### ERB: Behavioral Intoxication Rating Scale

The 6-point behavioral intoxication rating scale was conducted as previously described^23^. Briefly, animals were scored by two researchers according to the following behavioral scale: (1) no sign of intoxication; (2) hypoactivity; (3) slight intoxication (ataxia; slight motor impairment); (4) moderate intoxication (obvious motor impairment; dragging abdomen); (5) high intoxication (dragging abdomen; LORR); (6) extreme intoxication (LORR; loss of eye blink response). The behavioral intoxication rating scale was conducted at baseline as well as 15 min after each ethanol dose for a total of five assessments.

### ERB: Body Temperature

Body temperature was assessed using a Thermalert clinical monitoring thermometer (Physitemp, Clifton, NJ) with an electric thermometer probe inserted approximately 3 mm into the rectum and left in place for ≥45 s until a stable reading was obtained. Body temperature was assessed at baseline and again following each ethanol dose following completion of the behavioral intoxication rating scale for a total of five assessments. Difference (Δ) in body temperature as a consequence of cumulative ethanol dosing was calculated by subtracting body temperature following each ethanol dose from baseline body temperature. Room temperature was monitored daily and averaged 23°C.

### ERB: Blood Collection

Submandibular blood was collected at baseline and 15 min after each administration of ethanol following body temperature assessment to determine BECs using a GM7 Analyzer (Analox; London, United Kingdom).

### ERB: Tilting Plane

The tilting plane apparatus consisted of a clear Plexiglas box (18 cm × 7.5 cm × 9.5 cm) attached with a hinge to a frame with a glass panel floor. The box was tilted via an additional hinge attached to the base and a digital angle gauge was used to measure the angle at which subjects began to slide down the glass floor panel. At the time of testing, the mouse was placed on the apparatus facing away from the tilting hinge and the panel lifted slowly until the subject began to slide down the floor of the apparatus. The angle at which the mouse began to slide was measured and the procedure repeated for three consecutive trials per session. The three trials within each session were averaged and difference (Δ) in angle of slide as a consequence of cumulative ethanol dosing was calculated by subtracting angle of slide from the averaged baseline angle of slide from each ethanol dose angle of slide.

### ERB: Loss of Righting Reflex

Loss of righting reflex was assessed following the final dose of ethanol after completion of the tilting plane. Mice were placed on their back in a V-shaped trough and assessed for righting reflex. Loss of righting reflex in the ERB was defined as the inability of the mouse to right itself onto all four paws within 60 seconds.

### Pavlovian Ethanol Discrimination

The apparatus and sucrose access training used were as previously described^24^. Briefly, one side of the chamber had two cue lights that were located on either side of a liquid dipper receptacle. When activated, the dipper was raised for 4 seconds and presented 0.1 mL of sucrose (26% w/v). Head entries into the liquid receptacle were measured by infrared photobeam detectors. Additionally, general locomotor activity (number of beam breaks) was measured during the sessions with infrared photobeams that divided the chamber into four parallel zones. Acquisition Training was similar to previously described work^24^. Specifically, training sessions were conducted 5 days per week (M-F). The start of each training session begins with administration of ethanol (2 g/kg, i.g.) or water (i.g.) depending on the session, immediately after which rats are placed in the chambers for a 20 min timeout period. During this time no cue lights illuminated, no sucrose presented, head entries into the liquid receptacle not recorded. The 15-min session began after this delay. During ethanol training sessions, 10 cue light presentations occurred at random intervals for 15 sec. The offset of each of the 15-second cue light presentations was followed by presentation of sucrose. During water training sessions, no sucrose was delivered following the offset of the 10 cue light presentations. Ethanol and water training sessions varied on a double alternation schedule (E, E, W, W …). Training sessions were conducted until meeting criteria for discrimination: the mean of the first discrimination score from the preceding two ethanol sessions had to be greater than 3 of the mean of the first discrimination score for the preceding two water sessions^25,26^. Testing started when this criterion was met. The discrimination score was calculated as the number of head entries into the liquid receptacle during the 15-s light presentation minus the number of head entries during the 15-s immediately preceding the light presentation. This is an index of the conditioned response to the light cue. This score served as a measure of behavioral activation in response to the cue and under these training conditions, head entries during the light presentations increase on alcohol, but not water sessions. As such, the discrimination score is an index of the rat’s sensitivity to the interoceptive effects of ethanol. Importantly, only the first discrimination score (i.e., in response to the first light presentation) was used as this was prior to feedback from sucrose delivery. Locomotion during the test session (2-min) was recorded as beam breaks per min.

Test sessions were the same as training sessions except that they were 2 min in duration (after the 20 min delay) and consisted of a single light presentation. No sucrose was delivered following the offset of the light presentation so as not to influence subsequent behavior. In order to test whether SBI-553 would produce alcohol-like interoceptive effects or alter the interoceptive effects of alcohol, rats underwent two consecutive tests on a day. This strategy also maximized efficiency as testing SBI-553 pretreatment before water and ethanol could be tested in a single day. For the first test, rats were administered a dose of SBI-553 (0, 2, 6, 12 mg/kg, i.p.) then 10 min later, water administration (i.g.) and then placed in the chamber for the start of the test session (light presentation after the 20-min delay). Therefore, the pretreatment time from SBI-553 administration to testing interoceptive sensitivity was 30-min which is consistent with other assessments. Immediately after the first test session, rats were administered the ethanol training dose (2 g/kg, i.g.) followed immediately by the start of another test session (again, after the 20-min time out period). As such, the effects of SBI-553 were evaluated following both water and alcohol administration. All rats received all doses of SBI-553 over the course of 4 test sessions. Doses were counterbalanced across the 4 test sessions, and within a day the water test session preceded the ethanol test session. There was at least one intervening training session between each test session. Two male rats and one female rat died from gavage during testing, and as such data for 2 of the 4 doses are missing for these 3 rats.

### Data Analysis and Statistics

Mouse studies - Sample sizes were determined based on prior experiments in our lab. Graphpad Prism 10 was used for statistical analyses. Most mouse data was analyzed using a 2-Way ANOVA or 2-Way RM ANOVA to account for sex and treatment. If a main effect was present, Tukey’s multiple comparisons test was used for post-hoc comparisons. For ERB data, a 3-Way ANOVA was used to account for sex, treatment, and alcohol dose. If a main effect was present, Tukey’s multiple comparisons test was used for post-hoc comparisons. Mouse drinking data was analyzed using a 2-Way ANOVA to account for sex and treatment. If a main effect was present, a Fisher’s LSD test was used for post-hoc comparisons as male and female differences are well known in alcohol consumption^27^. For all data α≤0.05.

Rat study - A mixed-effects analysis with SBI-553 dose (4) and alcohol pretreatment (2) as within-subjects variables was used to test for the effects of SBI-553 on discrimination score and locomotion (due to missing values with attrition of 3 rats, see above). Because the training procedure purposefully produces differences in discrimination score between ethanol and water sessions (see criteria for testing), we *a priori* hypothesized a main effect of ethanol on discrimination score. \ Sidak’s multiple comparisons test was used for post-hoc analyses. For all data α≤0.05.

## Results

### SBI-553 reduces binge ethanol drinking and not sucrose drinking

In order to investigate the effect of SBI-553 on fluid consumption, we first performed a single-bottle DID paradigm (Fig. 1)^21^. We found that SBI-553 (12mg/kg, i.p.) significantly reduced the amount of ethanol consumed (g/kg) in the 2-hour drinking window (Fig. 1A) on the test day (day 4)^19^. (2-Way ANOVA. Main effect of sex F_(1, 33)_=4.824, p=0.0352, main effect of treatment F_(1, 33)_=5.356, p=0.027. Male vehicle n=9, male SBI-553 n=9. Female vehicle n=10, female SBI-553 n=9.). We saw no differences in sucrose consumption (Fig. 1B) in a separate, one bottle DID experiment in a different cohort of mice (2-Way ANOVA. Sex x treatment F_(1, 37)_=0.2036, p=0.6545. Male vehicle n=10, male SBI-553 n=10. Female vehicle n=10, female SBI-553 n=11.). To examine if SBI-553 would alter the preference for ethanol, we repeated the DID paradigm in a new cohort of mice with two bottles (Fig. 1C-E), one with 20% ethanol and one with water (counterbalanced, Male vehicle n=10, male SBI-553 n=10. Female vehicle n=10, female SBI-553 n=8.). Similar to the one bottle DID, we found a main effect of treatment group (2-Way ANOVA F_(1, 34)_=6.580, p=0.0149), surprisingly however there was not a main effect of sex (2-Way ANOVA F_(1, 34)_=0.4618, p=0.5014). Post-hoc analysis demonstrated that in the two-bottle choice experiment, SBI-553-mediated reduction in ethanol consumption is primarily driven by female mice (Fisher’s LSD. Female vehicle vs. SBI-553 p=0.0154). Examining preference for the ethanol bottle (Fig. 1E), there was a main effect of sex (2-Way ANOVA F_(1, 33)_=5.828, p=0.0215), but only a trend for an effect in treatment group (2-Way ANOVA F_(1, 33)_=3.153, p=0.0850). Post-hoc analysis demonstrated significant differences between vehicle vs. SBI-553 treated females (Fisher’s LSD p=0.0245) and between sexes in the SBI-553 treated animals (Fisher’s LSD p=0.0084). Interestingly, there was no difference in ethanol preference between the males treated with vehicle or SBI-553 (Fisher’s LSD p=0.9062). Additionally, we saw no difference in total fluid consumption (water and ethanol combined) in the two-bottle choice DID experiment (Fig. 1D) (2-Way ANOVA. Sex x treatment F_(1, 33)_ = 0.1814, p=.6730). These data suggest that SBI-553’s effects on fluid consumption may be specific to the reinforcing properties of ethanol.

**Figure 1.**
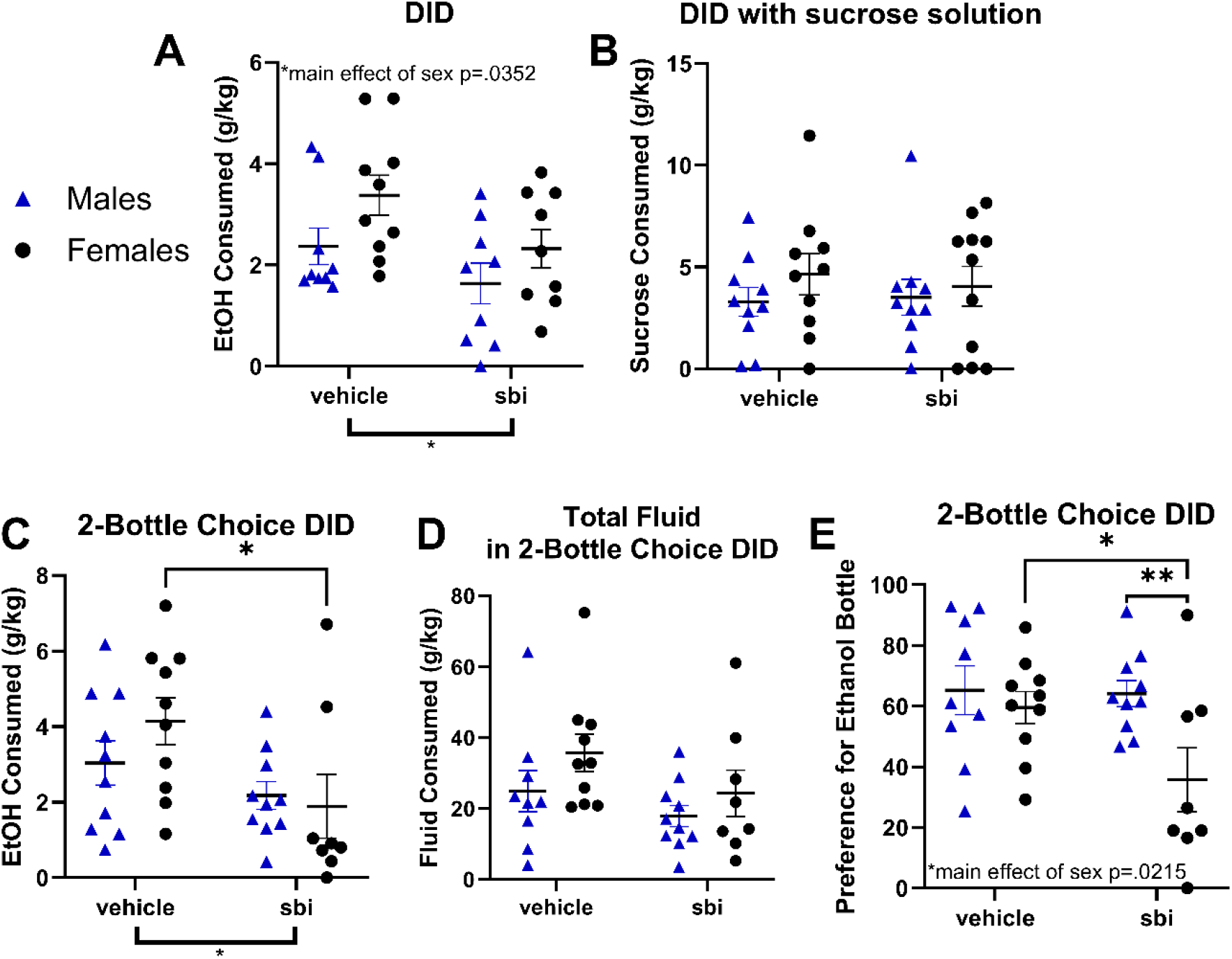
SBI-553 reduces binge-like ethanol drinking in mice. **A)** Ethanol consumed during DID normalized to body weight. **B)** Sucrose consumed during DID normalized to body weight. **C)** Ethanol consumed during 2-bottle choice DID normalized to body weight. **D)** The amount of water and ethanol combined consumed during 2-bottle choice DID. **E)** Preference for the ethanol bottle during 2-bottle choice DID.

### SBI-553 acutely suppresses locomotion in a novel context but does not affect avoidance behavior

In order to fully assess the therapeutic potential for SBI-553 we used a series of behavioral assays to determine if certain behaviors were affected outside of the context of ethanol (Fig. 2). We observed a small SBI-553 induced reduction in locomotion in the open field assay (Fig. 2A, B, male vehicle n=9, male SBI-553 n=9. Female vehicle n=9, female SBI-553 n=8). We observed this effect in both distance traveled (Fig. 2A) (2-Way ANOVA. Main effect of sex F_(1, 31)_=10.35, p=0.0030, main effect of treatment F_(1, 31)_=14.52, p=0.0006. Tukey’s multiple comparisons test: male treatment p=0.0148, SBI-553 male versus female p=0.0493) and velocity (Fig. 2B,2-Way ANOVA. Main effect of sex F_(1, 31)_=10.47, p=0.0029, main effect of treatment F_(1, 31)_=8.825, p=0.0057.). We saw no significant differences in avoidance behavior with regard to time spent in the center of the open field (Fig. 2C, male vehicle n=9, male SBI-553 n=9, female vehicle n=9, female SBI-553 n=8) of the open field (2-Way ANOVA sex x treatment F_(1, 31)_=0.0083, p=0.9281), time spent in the open arms (Fig. 2D) of the EPM (Male vehicle n=10, male SBI-553 n=10. Female vehicle n=10, female SBI-553 n=10) (2-Way ANOVA, sex x treatment F_(1, 36)_=0.6953, p=0.4099), or entries to the open arms (Fig. 2E) of the EPM (2-Way ANOVA, sex x treatment F_(1, 36)_=8.825e-030, p>0.9999).

**Figure 2.**
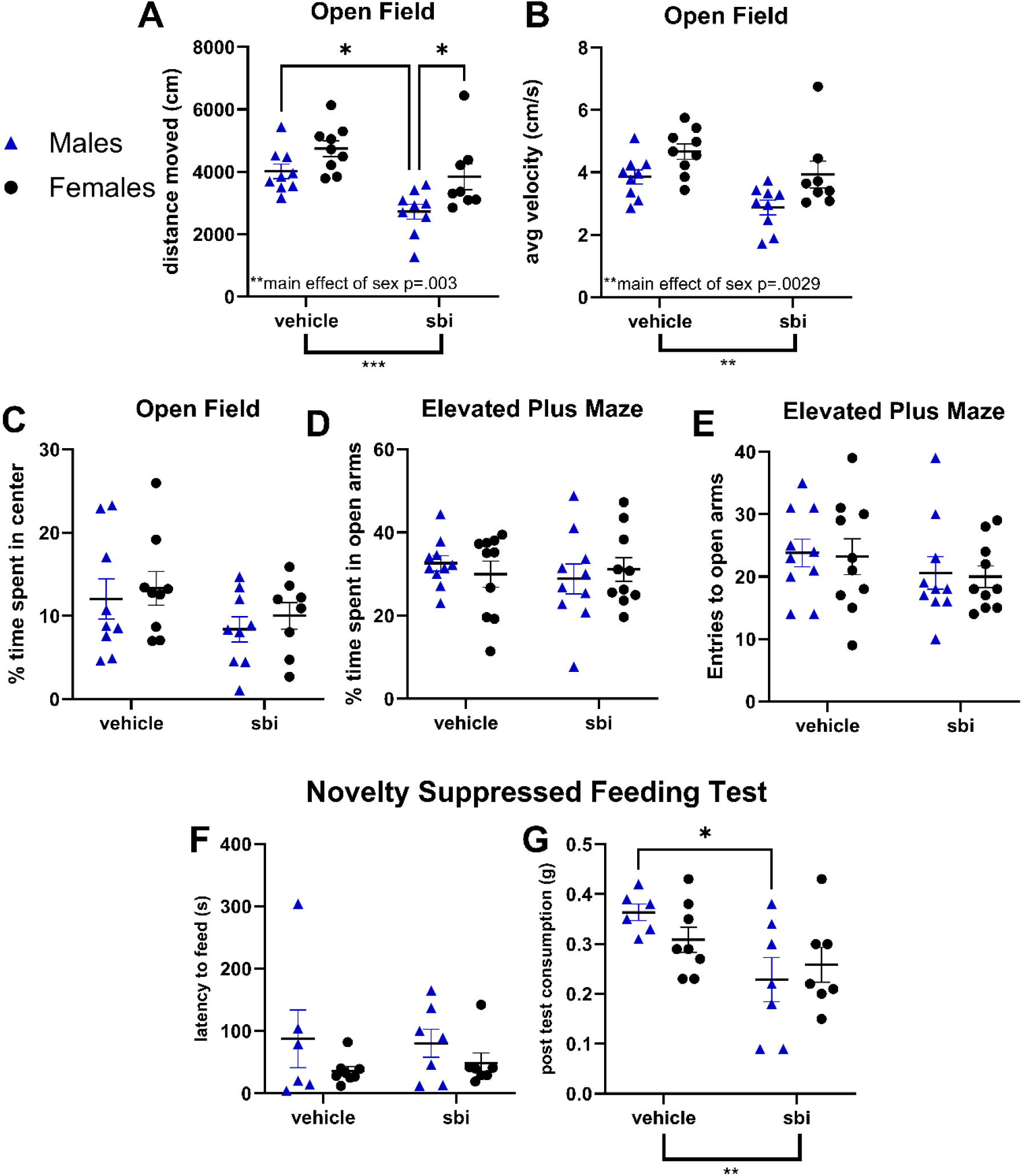
SBI-553 suppresses locomotion in a novel context but does not alter avoidance. **A)** Distance traveled in a 30-min open field assay. **B)** Average velocity in a 30-min open field assay. **C)** Time spent in the center of a 30-min open field assay. **D)** Time spent in the open arms of the elevated plus maze. **E)** Entries to the open arms of the elevated plus maze. **F)** Latency to feed in the novelty suppressed feeding test. **G)** Consumption in the home cage in the 10-min window following the novelty suppressed feeding test.

### SBI-553 reduces only post-test consumption in the novelty-suppressed feeding test (NSFT) and does not affect novel object recognition

Due to the involvement of Nts and Nts systems in consummatory and motivated behaviors, we next wanted to probe SBI-553’s action on latency to feed (amle vehicle n=6, male SBI-553 n=7, female vehicle n=8, female SBI-553 n=7) in the NSFT (Fig. 2F, G)^7,16,17,28–33^. We found no differences between treatment groups in the latency to feed (Fig. 2F) on a familiar palatable food in the NSFT (2-Way ANOVA, sex x treatment F_(1, 24)_=0.1736, p=0.6807). However, we did observe a significant reduction in the amount of food consumed in home cage palatable feeding during the 10-minute window (Fig. G) after the test (2-Way ANOVA, main effect of treatment F_(1, 24)_=8.006, p=0.0093, Tukey’s multiple comparisons test: male vehicle vs. SBI-553 p=0.0442). These results suggest a role for the NTSR1 system in motivation to feed on palatable foods depending on the hunger state of the mouse. The novel object recognition test (Fig. S1) can provide insight into how a treatment may affect memory performance and object recognition/object avoidance, which could potentially limit therapeutic efficacy in a clinical setting for a drug like SBI-553. We observed no significant differences in novel object preference (Fig. S1A) (Male vehicle n=6, male SBI-553 n=7. Female vehicle n=5, female SBI-553 n=6, 2-Way ANOVA Sex x treatment F_(1, 20)_=0.2103, p=0.6515) or in the latency to reach the 20s interaction criteria (Fig. S1B) between treatment groups (2-Way ANOVA Sex x treatment F_(1, 20)_=0.0019, p=0.9656). Together, these data suggest that avoidance behavior and cognition are unaffected by SBI-553 treatment, but that SBI-553 may modulate fasted-state consummatory behavior.

### SBI-553 does not alter avoidance behavior with systemic ethanol administration

In order to further assess the therapeutic potential for SBI-553, we repeated avoidance behavior measures in a separate group of animals that had a systemic injection of 1g/kg ethanol, generally considered an anxiolytic dose, prior to the assays (Fig. 3)^34^. This allowed us to measure avoidance behavior during ethanol intoxication in the context of SBI-553 treatment, which could potentially affect adherence to treatment in a clinical setting should SBI-553 and ethanol coadministration induce negative valence states. We found that ethanol pretreatment resulted in a significant main effect of sex in both distance traveled (Fig. 3A) (2-Way ANOVA, main effect of sex F_(1, 34)_=8.210, p=0.0071) and average velocity (Fig. 3B) (2-Way ANOVA, main effect of sex F_(1, 34)_=8.736, p=0.0056) in a novel open field (male vehicle n=9, male SBI-553 n=10, female vehicle n=9, female SBI-553 n=10). We observed no significant differences in time spent in the center of the open field (Fig. 3C, 2-Way ANOVA, sex x treatment F_(1, 34)_=1.950, p=0.1716), time spent in the open arms of an EPM (Fig. 3D, 2-Way ANOVA, sex x treatment F_(1, 31)_=0.8700, p=0.3582), or entries to the open arms of an EPM (Fig. 3E, 2-Way ANOVA, sex x treatment F_(1, 31)_=0.6589, p=0.4231, EPM: Male vehicle n=9, male SBI-553 n=8, female vehicle n=9, female SBI-553 n=9). These data largely show that avoidance behavior is not impacted by SBI-553 treatment with exposure to systemic ethanol exposure.

**Figure 3.**
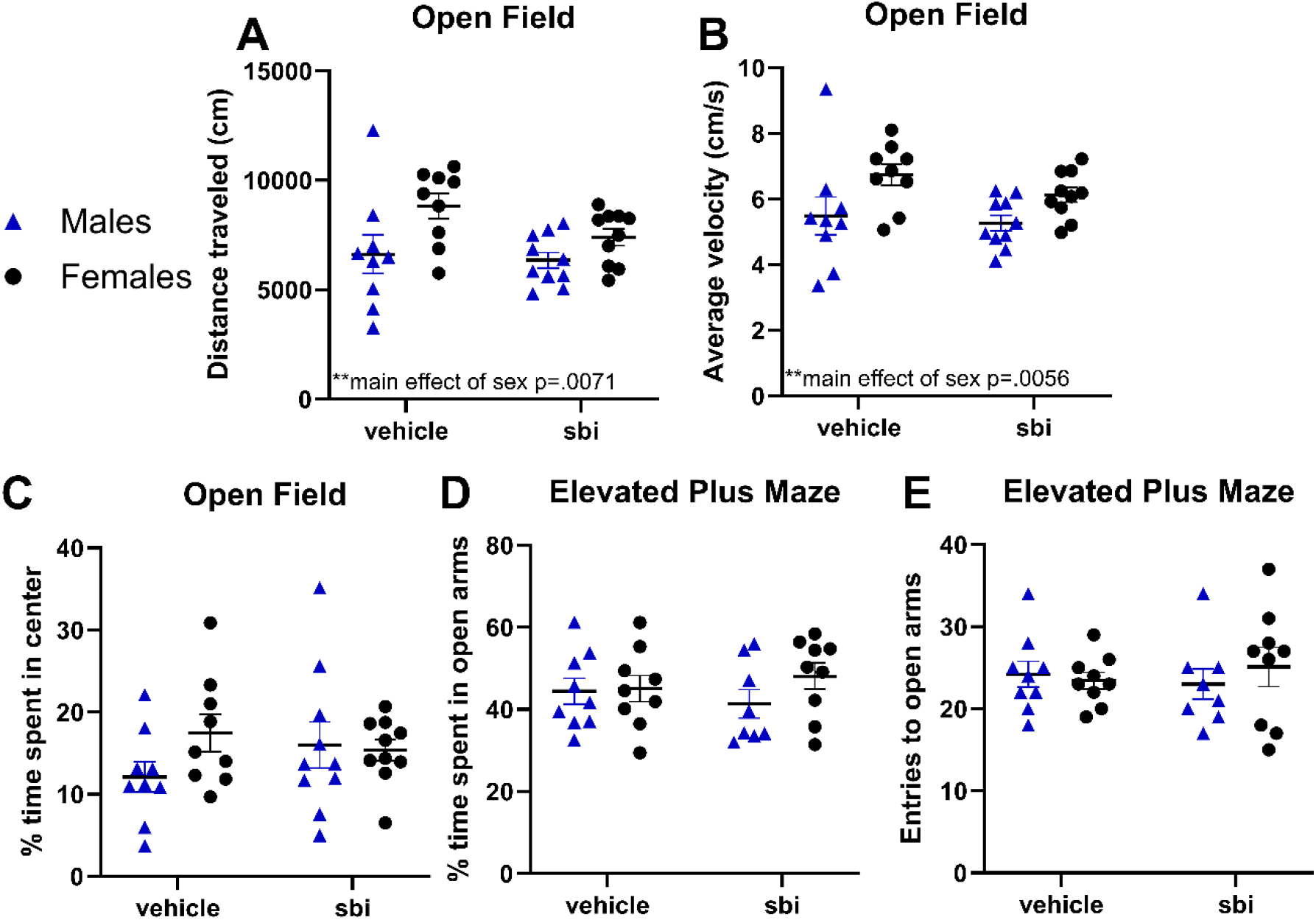
SBI-553 does not alter locomotion or avoidance with ethanol on board. **A)** Distance traveled in a 30-min open field assay. **B)** Average velocity in a 30-min open field assay. **C)** Time spent in the center of a 30-min open field assay. **D)** Time spent in the open arms of the elevated plus maze. **E)** Entries to the open arms of the elevated plus maze.

### SBI-553 modulates physiological response to ethanol in the Ethanol Response Battery

In order to assess physiological responses to ethanol in mice we used the ERB procedure (Fig. 4) described in the methods section (male vehicle n=10, SBI-553 sbi n=10, female vehicle n=10, female SBI-553 n=10). In this experiment, we found that SBI-553 reduces ethanol intoxication score. Although these data did not survive post-hoc comparison, examination of the data demonstrates that the effects are driven by the SBI-553 treated female mice. (Fig. 4A, 3-Way ANOVA, main effect of treatment F_(1, 36)_=10.33, p=0.0028, main effect of sex F_(1, 36)_=5.688, p=0.0225). We also observed significant ethanol dose x treatment, ethanol dose x sex, and treatment x sex interactions (ethanol dose x treatment F_(3, 108)_=6.894, p=0.0003. Ethanol dose x sex F_(3, 108)_=5.131, p=0.0023. Treatment x sex F_(1, 36)_=4.742, p=0.0361). Examining ethanol-induced hypothermia, (Fig. 4B) we found a main effect of sex on change in body temperature that is again driven by SBI-553-treated females based on a significant treatment x sex interaction, and a significant ethanol dose x treatment x sex interaction (3-Way ANOVA, ain effect treatment F_(1, 36)_=10.33, p=0.0028, sex F_(1, 36)_=5.688, p=0.0225. Interactions: EtOH dose x treatment F_(3, 108)_=6.894, p=0.0003, EtOH dose x sex F_(3, 108)_=5.131, p=0.0023, treatment x sex F_(1, 36)_=4.742, p=0.0361, EtOH dose x treatment x sex F_(3, 108)_=3.436, p=0.0195). Investigating ethanol metabolism by measuring BECs, (Fig. 4C) we examined striking, sex-dependent effects of SBI-553 with main effects of sex and significant ethanol dose x sex, treatment x sex, and ethanol dose x treatment x sex interactions (3-Way ANOVA, main effect of sex F_(1, 36)_=17.51, p=0.0002. Interactions: EtOH dose x sex F_(3, 108)_=8.333, p<0.0001, treatment x sex F_(1, 36)_=9.069, p=0.0047, EtOH dose x treatment x sex F_(3, 108)_=3.761, p=0.0130.). We also observed post-hoc differences in several male treatment groups (noted by +) and differences between males and females in the SBI-553 group (noted by $) in BECs at all ethanol doses observed in the study (3-Way ANOVA, Tukey’s: 0.5 M vehicle vs. treatment p<.0001, 0.5 M vs. F SBI-553 p<.0001, 1.0 M tx p=.0032, 1.0 M vs F sbi p<.0001, 2.0 M tx p=.0254, 2.0 M vs F sbi p<.0001, 3.0 MvsF sbi p=.0063). Finally, in the tilting plane assay (Fig. 4D) we saw a significant main effect of SBI-553 treatment on ethanol-induced motor impairment based on change in angle of slide from baseline (3-Way ANOVA, main effect treatment F_(1, 36)_=19.41, p<0.0001.). We also observed significant treatment x sex, and ethanol dose x treatment interactions (3-Way ANOVA, interactions: EtOH dose x treatment F_(3, 108)_=5.166, p=0.0022, treatment x sex F_(1, 36)_=5.188, p=0.0288). Our data also shows that male mice exhibit a more robust attenuation of ethanol-induced motor impairment with significant post-hoc test results (3-Way ANOVA, Tukey’s multiple comparisons test: 1.0g/kg etoh male vehicle vs. SBI-553 p=0.0063, 2.0g/kg EtOH male vehicle vs. SBI-553 p=0.0229).

**Figure 4.**
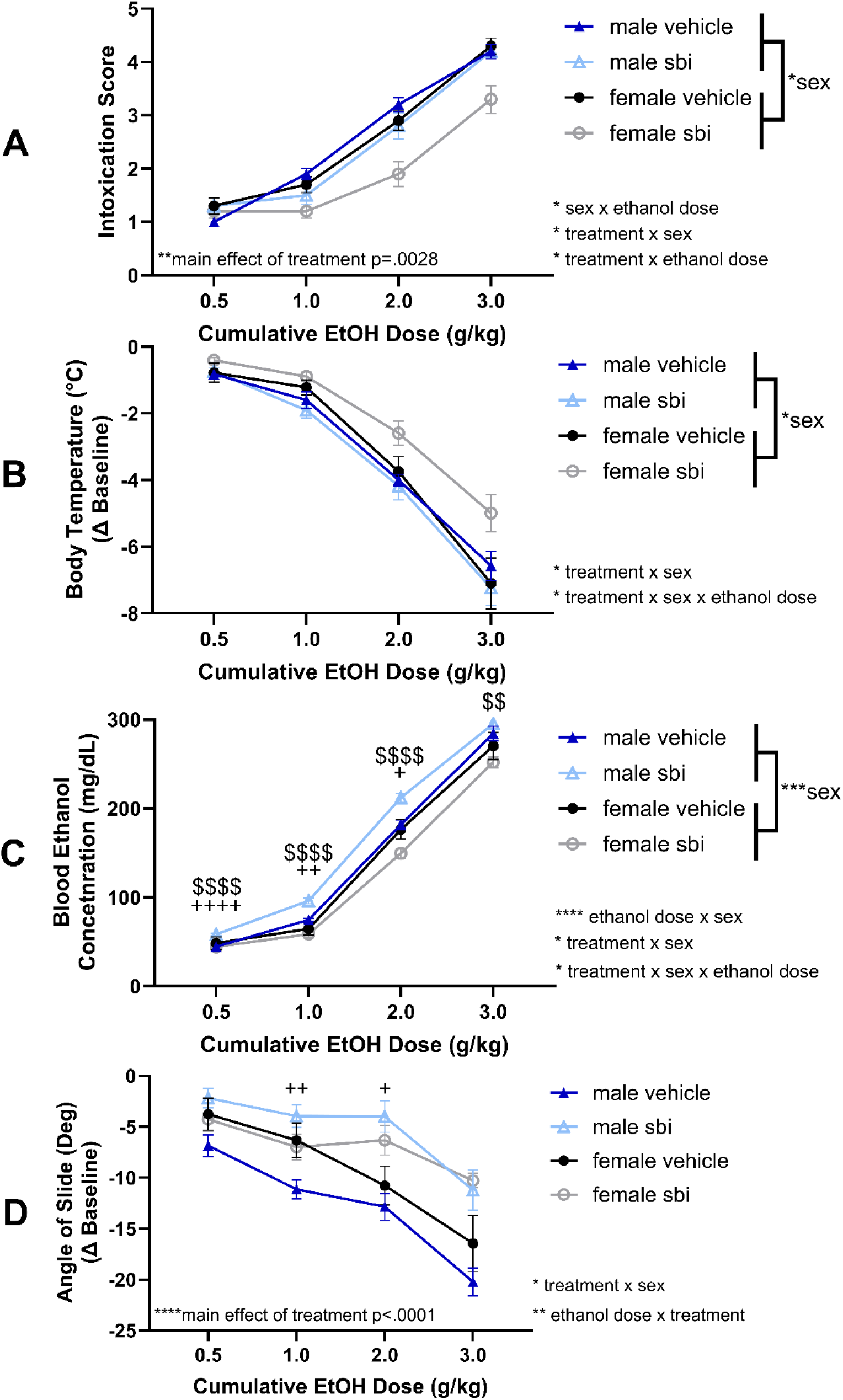
SBI-553 modulates physiological response to ethanol. $=significant sex difference in sbi group. +=significant difference between treatment groups in males. **A)** Ethanol intoxication scores at increasing cumulative doses of ethanol administered during the ethanol response battery. **B)** Change in body temperature from baseline. **C)** Blood ethanol concentrations. **D)** Change in angle of slide from baseline in the tilting plane assay.

### SBI-553 reduces interoceptive sensitivity to ethanol in rats

Because our data in mice suggested that there were differences in ethanol consumption, preference, physiological response, and intoxication following SBI-553 pretreatment, we hypothesized that SBI-553 may alter the interoceptive qualities of ethanol. To examine this, we used a Pavlovian model of drug discrimination (Fig. 5) in male and female rats and tested 3 doses of SBI-553 (2, 6, and 12 mg/kg, i.p.). Males and females were analyzed separately. Examining discrimination scores in male rats (Fig. 5A), we found a main effect of SBI-553 (F_(3, 21)_=5.768, p=0.0049). Post-hoc tests showed an effect of ethanol in the male vehicle group (p00053), confirming stimulus control, but this difference was not observed in any SBI-553 treatment groups. SBI-553 abolished the difference (ethanol versus water) at all doses (2, 6, 12mg/kg). There was no significant effect of ethanol on locomotion (Fig. 5B), but there was a main effect of SBI-553 (F_(3, 18)_=5.610, p=0.0068) and no interaction effect. Unfortunately, one of the chambers had a malfunctioning infrared detector for locomotor activity, and thus data from one of the rats is not included in this analysis. This significant increase in locomotor behavior was surprising, and lead us to interpret that the reduction in discrimination score was not related to a potential nonspecific decrease in locomotor behavior.

**Figure 5.**
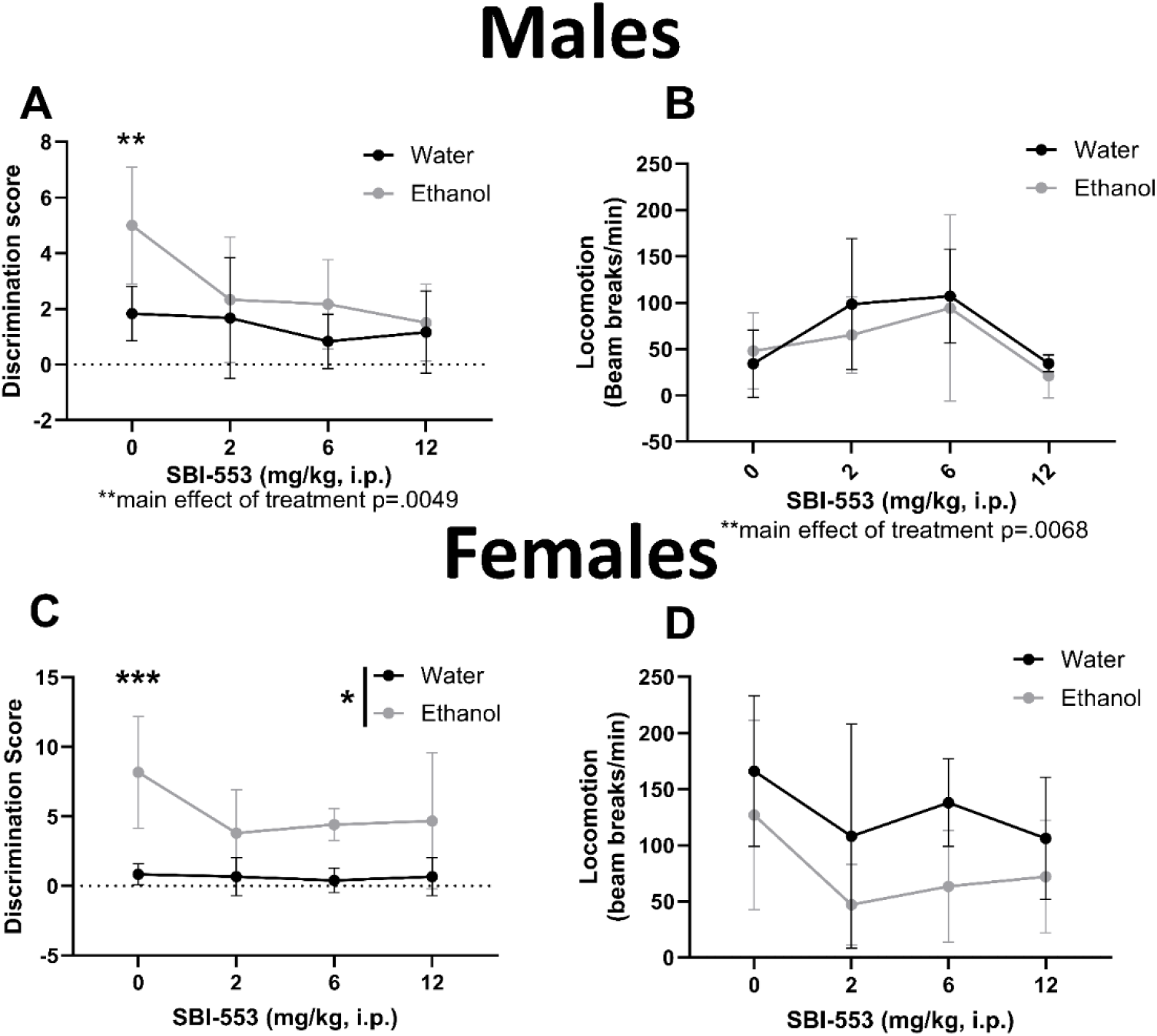
SBI-553 reduces the interoceptive effects of ethanol in rats. **A)** Male discrimination scores at different doses of SBI-553 during test sessions. **B)** General locomotor behavior in males during the SBI-553 test sessions. **C)** Female discrimination scores at different doses of SBI-553 during test sessions. **D)** General locomotor behavior in females during the SBI-553 test sessions.

In female rats (Fig. 5C), there was a main effect of ethanol (F_(1, 5)_=15.59, p=0.0109) on discrimination score, with higher discrimination scores following ethanol treatment. Post-hoc tests showed an effect of ethanol in the female vehicle group (p=0.0008), confirming stimulus control, but not in any SBI-553 treated groups. Although, there were trends for higher discrimination scores on the alcohol sessions relative to water at the 6 and 12 mg/kg doses (p<0.07). Therefore, SBI-553 abolished the difference between ethanol versus water, indicating a blunting of the interoceptive effects of ethanol). There was no significant main effect of ethanol, SBI-553, nor a significant interaction effect when locomotion was examined (Fig. 5D). Together these data suggest that SBI-553 blunted the interoceptive effects of alcohol in both male and female rats.

## Discussion

### AUD

Alcohol use is prevalent within the population with over 85% of adults in the United States reporting consuming alcohol in their lifetime, and over 25% of adults in 2019 reporting alcohol misuse (binge drinking) in the past month^35^. The ongoing global COVID-19 pandemic has only contributed to an increase in alcohol consumption and binge drinking^36^. Although these statistics are staggering, there are currently only 3 pharmacotherapeutic treatment strategies that are FDA approved for AUD, disulfiram, naltrexone, and acamprosate; and, less than 9% of the persons who could be aided by these medications receive them as part of their treatment^36,37^. This further exemplifies the need to explore novel treatment strategies, and untapped receptor systems, to combat alcohol misuse and increase harm reduction strategies.

### SBI-553 does not alter avoidance behavior

If SBI-553, or similar compounds, can be considered as a medication to treat AUD, it is important to assess how it may otherwise alter behavior on the affective spectrum and cognitive behaviors. Undesirable side effects in affect and cognition would likely decrease medication compliance, and probably contribute to the low use of the 3 FDA approved medications listed above. To begin to probe these concepts, we investigated if SBI-553 would alter avoidance behavior by examining behavior in the open field, EPM and NSFT. Additionally, we probed how a 1g/kg, “anxiolytic dose” of ethanol could impact these behaviors, as neurotensin and ethanol have previously been demonstrated to cross sensitize^10^. Further, we investigated both novel object recognition and the time it took to engage 20 seconds with a novel object. Our data suggests that SBI-553 did not impact avoidance behavior (with or without the 1 g/kg dose of ethanol) or memory performance. There were, however, a few differences observed. In a novel open field, SBI-553 slightly but significantly reduced both the velocity and the distance traveled. This was in contrast to previous reports where SBI-553 did not alter locomotor behavior in a familiar open field context, alter a wire grasping task, or cause sedation. We believe that our observations are due to the novelty of the open field, and that SBI-553 reduced novelty induced hyperlocomotion. Further, we observed no effect of SBI-553 on locomotor activity in the novel open field with 1g/kg ethanol delivered systemically prior to the test, suggesting that unlike classical agonists of NTSR1, SBI-553 does not cross sensitize with ethanol^10^. Interestingly, we also found that SBI-553 pretreatment reduced food consumption in the home cage following the NSFT. This result was driven by the male animals and suggests that SBI-553 may alter food consumption under distinct motivation states (see below for sucrose consumption discussion.) These data are critical as Nts negatively regulates food consumption in the central nervous system, but in the periphery it can promote fat absorption and weight gain (for detailed discussion, see Ramirez-Virella & Leinninger 2021)^16^.

### Modulation of ethanol consumption by SBI-553

Here we observe that SBI-553 reduces binge-like ethanol consumption in mice in two DID paradigms, single bottle and 2-bottle choice DID, both administered in the home cage. When investigating sucrose drinking (single bottle) in the DID paradigm, we observed no differences between vehicle or SBI-553 groups, suggesting that SBI-553’s effects on reinforcing fluid consumption could be specific to ethanol. This is supported by the fact that we did not observe differences in total fluid consumption (ethanol and water combined) between vehicle and SBI-553 groups in the 2-bottle choice DID experiment. Although we did observe a locomotor effect of SBI-553 in the novel open field, again, we believe that this did not contribute to reduced ethanol drinking, as we did not see differences in total fluid consumption in 2-bottle choice DID or sucrose drinking in DID in the home cage. Additionally, previous studies have not observed locomotor effects of SBI-553 in familiar contexts or different assays^19^. This suggests that the observed locomotor depression depends on the novelty of the open field arena, and that repeated locomotor studies would likely show no significant differences between treatment groups when the novelty of the environment is not a factor. Additionally, other work has shown that SBI-553 does not impact latency to grasp the wire in the wire hang assay, while traditional NTSR1 agonists significantly increase latency to grasp the wire, suggesting that SBI-553 does not affect other measures of motor impairment in which novelty is not implicated in the results of the assay^19^. Finally, in our rat study, we saw a surprising increase in locomotor behavior due to SBI-553 driven by the lower doses examined (2 and 6 mg/kg). In future studies it would be interesting to see if this effect was also observed in mice at lower doses.

### ERB and Pavlovian discrimination data suggest SBI-553 reduces sensitivity to ethanol

Researchers working with models of substance use and substance use disorders tend to focus on the brain for development of therapeutic strategies and target identification, but here our data suggests that actions of ethanol on the body as a whole could be a key component of regulating ethanol consumption. We observed that along with a reduction in ethanol consumption, SBI-553 affects the rodent’s physiological response to ethanol administration and the interoceptive effects of ethanol. This is a key finding since sensitivity to alcohol in clinical trials predicts therapeutic benefits of AUD pharmacotherapies^38^. This could explain the effect of SBI-553 on ethanol consumption compared to other fluids, including sucrose which is another caloric, rewarding fluid. SBI-553 reductions in alcohol consumption and sensitivity make it a promising pharmacotherapy for AUD. SBI-553 lowered observed intoxication levels and attenuated ethanol-induced hypothermia in female mice, which could be due to either neuronal mechanisms or peripheral mechanisms, however Nts is known to modulate body temperature peripherally via NTSR2^39–41^. Interestingly, male SBI-553 mice had a slight but significantly higher BEC than vehicle controls, but paradoxically showed reduced ethanol -induced motor impairment. These findings lead us to a hypothesis that SBI-553 reduces the interoceptive effects of ethanol, which reduces motivation for ethanol drinking. This assertion is supported by the rat study investigating Pavlovian ethanol discrimination, which showed that SBI-553 reduced sensitivity to the interoceptive effects of ethanol vis-a-vi lower ethanol discrimination scores when treated with SBI-553 compared to vehicle. If SBI-553 modulated reward in general, we would expect to see differences in sucrose consumption. Finally, we did observe a decrease in palatable food consumption in the home cage following the NSFT test. This test is conducted in a fasted state which alters motivation for the reward, whereas our sucrose DID test was not performed in a fasted state. It would be interesting in future studies to examine how an animal’s internal motivational state to seek rewards is modulated by SBI-553 and/or similar compounds.

### Sex as a biological variable

We observed many sex differences throughout the data discussed in this study. These differences observed highlight the importance of examining sex as a biological variable, especially given that there are sex and gender differences in alcohol use in humans^42^. That being said, these differences can be challenging to interpret, especially considering that female mice drink significantly more ethanol than male mice, which is the opposite of behavior observed in human populations (although these numbers are changing due to societal factors)^43–46^. Men drink an average of 19 liters of alcohol per year while women consume 6.7 liters per year on average^47^. Despite this data, normalization to body weight may make these consumption values more congruent. Plus, binge-drinking in particular, which is what we model in this study, is increasing in women at a faster rate than in men^43^. One source of the differences we observe here could be that Nts expression is increased in response to estrogen due to an estrogen response element in the gene^48^. Differences in the neurotensin system throughout the central nervous system could also be mediating some of these sex differences we observed. Female rats express around four times as much Nts in the anteroventral periventricular nucleus^49^. Additionally, manipulations of Nts neurons in the central amygdala, a brain region well-known to respond to and control ethanol consumption, modulate ethanol drinking in male mice, but not in female mice^2^. Ultimately, in this study we investigated systemic administration of SBI-553, which will likely exert diverse effects throughout the brain and body, making the source of the sex differences observed difficult to parse currently, however this is a promising area for future study.

### Study limitations

The data presented in this paper and by others point to the potential for β-arrestin positive/Gq negative bias at NTSR1 as a novel AUD/SUD pharmacotherapeutic that could reduce ethanol and drug consumption with limited side effects^19^. Additional research, however is needed in this space to fully assess the therapeutic potential for clinical settings. First, the work presented here only examines acute dosing of SBI-553. Observing maintained efficacy in reducing ethanol consumption in longer term models of ethanol consumption and repeated SBI-553 dosing across several weeks and/or months is critical to examine changes in behavioral and neural plasticity that may occur in AUD/SUD specific circuits. Given the roles of Nts in reward modulation, consummatory behavior, and key metabolic processes that regulate consumption, studying the effects of SBI-553 on diet preference and physical activity are particularly important^7,17^. Especially critical to consider is the fact that NTSR1 is most densely expressed in the gut where it regulates fat absorption postprandially^50,51^. How this impacts diet preference and regulation of body weight is important to consider, especially since other work has shown a role for NTSR1 in body weight and physical activity regulation in the ventral tegmental area^30,31,52^. Additionally, this work would be well-supported by an operant behavior study with SBI-553. In such a study, motivation for ethanol rewards could be assessed by using fixed ratio and progressive ratio reinforcement schedules in an operant behavior paradigm. Other studies have shown that SBI-553 reduces operant responding for psychostimulants and Nts-induced changes in VTA neuron activity and dopamine release^19,53^. Based on parallels in our data to the previous work by Slosky et al., we would expect a similar result in the context of ethanol. Finally, determining how SBI-553 modulates ethanol reward learning in the conditioned place preference assay would be useful. Given that SBI-553 reduces psychostimulant conditioned place preference in other work, we would expect a similar result with ethanol as the conditioned reinforcer^19^.

## Conclusion

Here, we present evidence that systemic SBI-553 modulates ethanol-related behaviors including a reduction in the amount of ethanol consumed in a binge-like ethanol drinking model (2-bottle choice and one bottle drinking in the dark, or DID) in mice. We find that SBI-553 does not modulate avoidance behaviors but does suppress locomotion in mice in a novel context. However, our sucrose drinking and two-bottle choice DID experiments suggest that the reduction in ethanol drinking that we see is not related to this locomotor effect of SBI-553. We find changes in the physiological response to ethanol in mice treated with SBI-553. Many of the effects in mice were sex specific. Lastly, we find that SBI-553 reduces ethanol discrimination score in rats, which we believe is a key finding supporting our hypothesis that SBI-553’s effects on ethanol drinking are primarily related to changes in interoceptive sensitivity to ethanol. These data suggest that allosterism at NTSR1 by β-arrestin-biased ligands lacking Gq activity may be a promising strategy for the treatment of AUD, and that SBI-553 could be a strong pharmacotherapeutic candidate for the treatment of AUD.

## Supplemental Figures

**Supplementary Figure 1.**
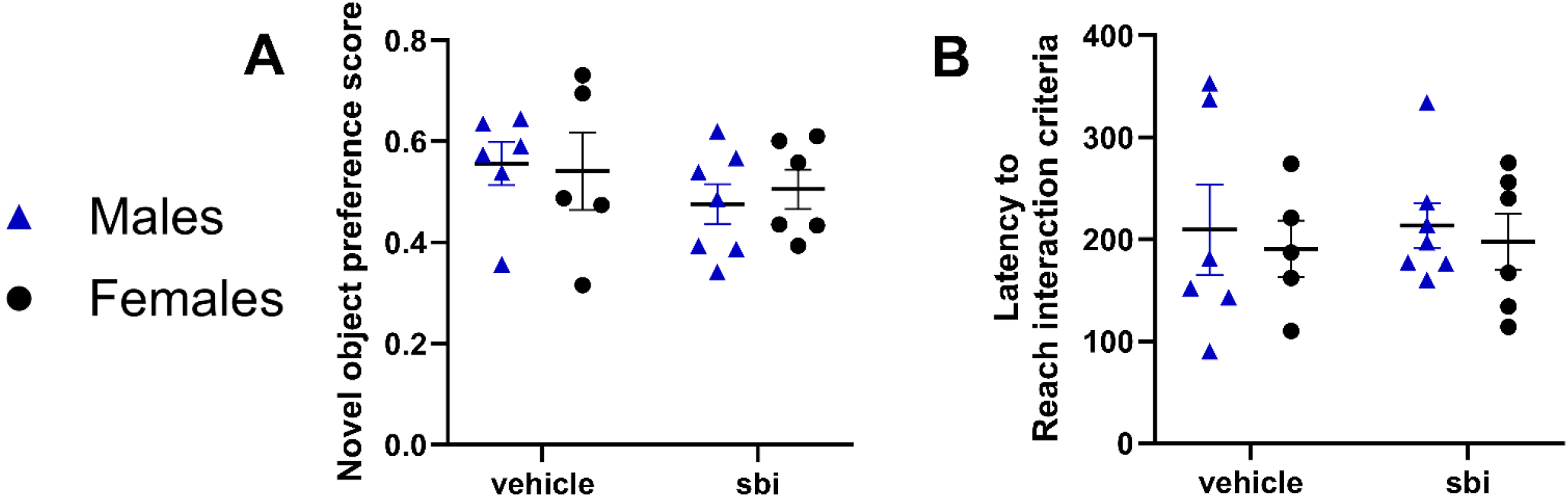
SBI-553 does not affect novel object recognition. **A)** Novel object preference scores. **B)** Latency to reach 20s of interaction combined between the novel and familiar object.

**Supplementary Figure 2.**
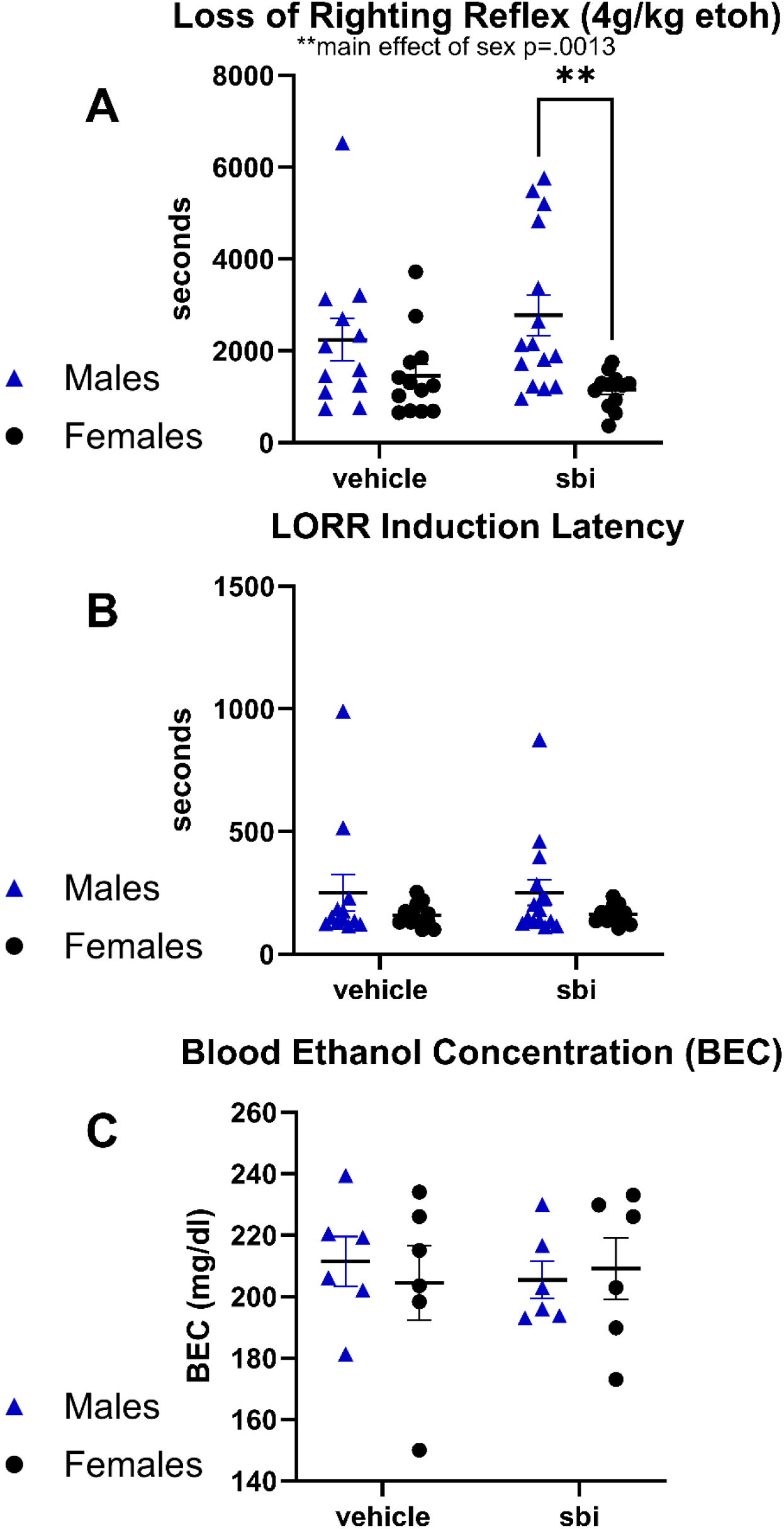
SBI-553 does not affect acute ethanol metabolism 15 minutes after a 2g/kg dose of ethanol or sedation by a 4g/kg dose of ethanol. **A)** Loss of righting reflex time after a 4g/kg dose of ethanol. **B)** Latency to lose righting reflex after a 4g/kg dose of ethanol. **C)** Blood ethanol concentration 15 min after 2g/kg ethanol injection based on analysis of plasma from trunk blood.

**Supplementary Figure 3.**
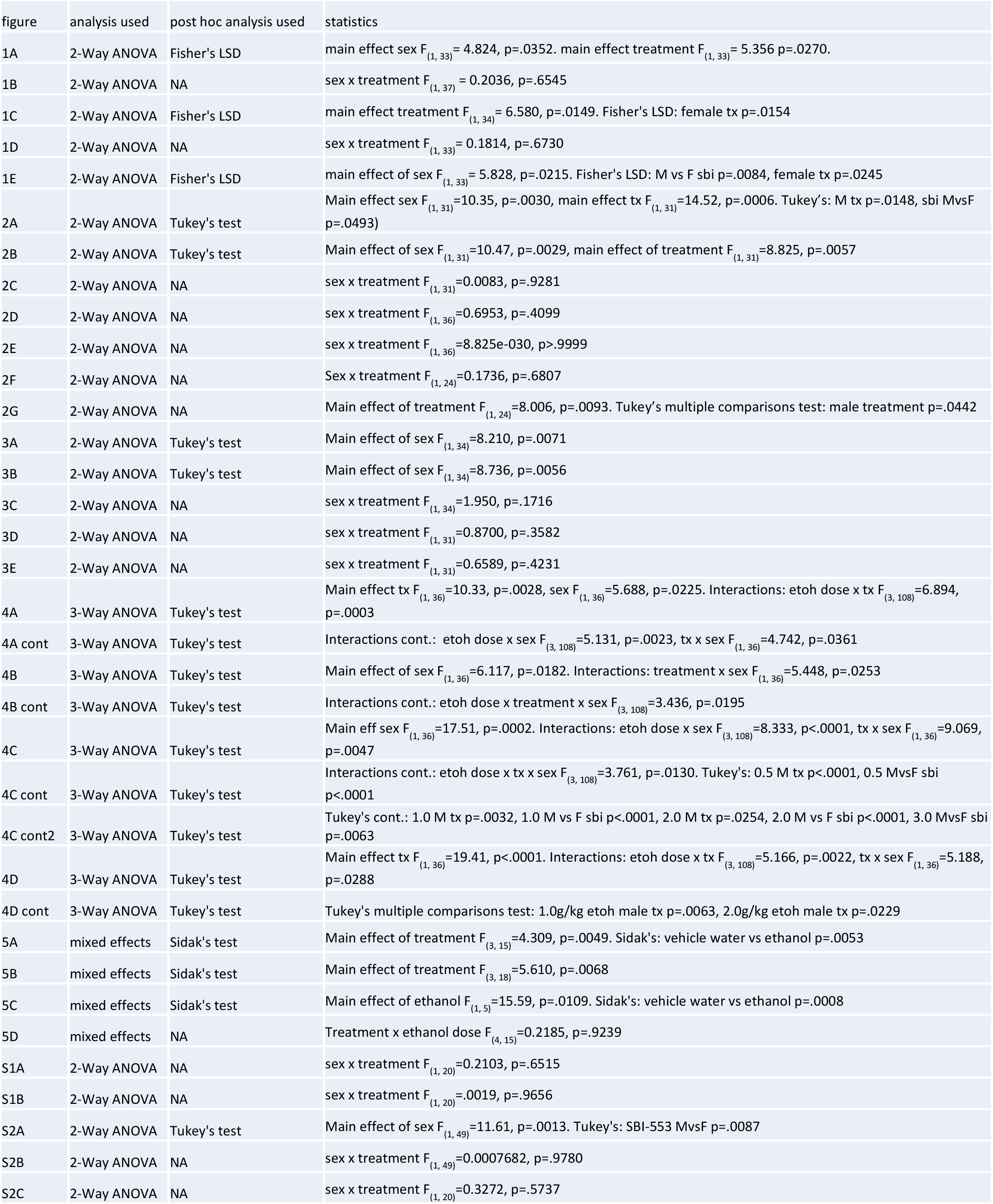
Data Table.

